# Single neuron contributions to the auditory brainstem EEG

**DOI:** 10.1101/2024.05.29.596509

**Authors:** Paula T. Kuokkanen, Ira Kraemer, Christine Koeppl, Catherine E. Carr, Richard Kempter

## Abstract

The auditory brainstem response (ABR) is an acoustically evoked EEG potential that is an important diagnostic tool for hearing loss, especially in newborns. The ABR originates from the response sequence of auditory nerve and brainstem nuclei, and a click-evoked ABR typically shows three positive peaks (‘waves’) within the first six milliseconds. However, an assignment of the waves of the ABR to specific sources is difficult, and a quantification of contributions to the ABR waves is not available. Here, we exploit the large size and physical separation of the barn owl first-order cochlear nucleus magnocellularis (NM) to estimate single-cell contributions to the ABR. We simultaneously recorded NM neurons’ spikes and the EEG in owls of both sexes, and found that ≳ 5, 000 spontaneous single-cell spikes are necessary to isolate a significant spike-triggered average response at the EEG electrode. An average single-neuron contribution to the ABR was predicted by convolving the spike-triggered average with the cell’s peri-stimulus time histogram. Amplitudes of predicted contributions of single NM cells typically reached 32.9 ± 1.1 nV (mean ± SE, range: 2.5 − 162.7 nV), or 0.07 ± 0.02% (median ± SE; range from 0.01% to 1%) of the ABR amplitude. The time of the predicted peak coincided best with the peak of the ABR wave II, independent of the click sound level. Our results suggest that individual neurons’ contributions to an EEG can vary widely, and that wave II of the ABR is shaped by NM units.

**Significance statement:** The auditory brainstem response (ABR) is a scalp potential used for the diagnosis of hearing loss, both clinically and in research. We investigated the contribution of single action potentials from auditory brainstem neurons to the ABR and provide direct evidence that action potentials recorded in a first order auditory nucleus, and their EEG contribution, coincide with wave II of the ABR. The study also shows that the contribution of single cells varies strongly across the population.

## Introduction

ABRs typically exhibit three early peaks, generated in the brainstem by local current sources arising from the auditory nerve as well as first- and second-order auditory nuclei in succession. These local current sources give rise to extracellular field potentials (EFPs) whose origins are not well understood, despite their clinical relevance. Studies of cortical pyramidal cells have led to the widespread assumption that EFPs have their origins mainly in synaptic dipoles (Eccles, 1951; Klee et al., 1965; Creutzfeldt et al., 1966a,b; Nunez and Srinivasan, 2006; da Silva, 2013; Ilmoniemi and Sarvas, 2019). However, other neuronal sources can also contribute, because the source of EFPs depends on the morphology of potential neuronal sources and synchrony of their activity (Gold et al., 2006; Kuokkanen et al., 2010; Lindén et al., 2011; McColgan et al., 2017; Rimehaug et al., 2023). Identifying the sources of brainstem EFPs, and their contributions to the ABR, should both inform models of the ABR and provide further insights into different types of hearing loss. We show here the contributions of single neurons to the ABR.

ABRs were first detected in the 1950s (Dawson, 1954; Geisler et al., 1958), and have been widely used in the clinic for decades (Geisler, 1960; Clark et al., 1961). Furthermore, ABRs are used in common basic hearing tests in animal research (e.g., Zheng et al., 1999; Akil et al., 2016; Kim et al., 2022). Models of the ABR (e.g. Melcher and Kiang, 1996; Ungan et al., 1997; Goksoy et al., 2005; Riedel and Kollmeier, 2006; Colburn et al., 2008; Verhulst et al., 2015, 2018) have helped to clarify ideas about its sources and its binaural components, but have remained difficult to validate experimentally (Riedel and Kollmeier, 2003; Tolnai and Klump, 2020). Most ABR models incorporate the *unitary response* (UR) (Melcher and Kiang, 1996; Dau, 2003; Schaette and McAlpine, 2011; Rønne et al., 2012; Verhulst et al., 2015, 2018), which is the expected average spike-triggered response at the EEG electrode related to the activation of a single neuronal source. The UR typically also includes the full structurally correlated cascade of activations in other brainstem nuclei. When convolved with the peri-stimulus time histogram of that (initial) source, the UR predicts the contribution of that source (and related later sources) to the ABR response. There are, however, many possible UR-solutions to a given ABR waveform, where each solution imposes a set of boundary conditions related to the source of the UR in the cell morphology. Furthermore, URs have been difficult to measure, leading to methods to estimate them indirectly for the whole brainstem by deconvolution from the ABR and models of firing rates (e.g. Elberling, 1978; Dau, 2003; Rønne et al., 2012). The deconvolution method is adequate for modeling expected ABR responses from various stimuli, but lacks precision about the sources whose activity might be correlated with changes in the UR.

To measure the contribution of individual nucleus magnocellularis (NM) neurons to the EEG, we took advantage of the large size and physical separation of the first-order auditory nuclei in birds (Kubke et al., 2004). NM units have high spontaneous firing rates (Köppl, 1997a), which allows recording of tens of thousands of spontaneous spikes; using spontaneous spikes minimizes stimulus-induced correlations in the EEG signal, and averaging over many spikes diminishes the noise at the EEG electrode to isolate the spike-triggered average of NM units. Convolving a measured PSTH with the spike-triggered average EEG allowed us to predict the NM neuron’s contribution to the ABR. Unlike the UR approach above, we specifically could exclude most structural correlations, and we only measured the UR for NM, which allowed us to dissect its contribution to the ABR.

## Materials and Methods

All the data analysis was done with Matlab 9.0 (version 2016a, MathWorks, Natick, MA). All the data was re-sampled to 50 000 Hz before analysis for consistency with previous analyses, for example in Kuokkanen et al. (2010, 2018).

### Experimental paradigm

The experiments were conducted in the Department of Biology of the University Maryland. Thirteen barn owls (Tyto furcata) of both sexes were used to collect the data at 27 EEG recording locations and for 151 intracranial recording locations. Many animals were studied in two or three separate physiology experiments, spaced approximately a week apart. Procedures conformed to NIH Guidelines for Animal Research and were approved by the Animal Care and Use Committee of the University of Maryland. Anaesthesia was induced by intramuscular injections of 16 mg/kg ketamine hydrochloride and 3 mg/kg xylazine. Similar supplementary doses were administered to maintain a suitable plane of anaesthesia. Body temperature was maintained at 39^◦^C by a feedback-controlled heating blanket. More details can be found in Carr et al. (2015).

#### Acoustic stimuli

Recordings were made in a double walled sound-attenuating chamber (IAC Acoustics, IL) with 65 − 75 dB noise reduction for 1 − 8 kHz. Acoustic stimuli were digitally generated by custom-made software (“Xdphys” written in Dr. M. Konishi’s lab at Caltech) driving a signal-processing board (DSP2 (Tucker-Davis Technologies (TDT), Gainesville, FL). Acoustic signals were calibrated individually at the start of each experiment, using built-in miniature microphones (EM3068; Knowles, Itasca, IL) inserted into the owl’s left and right ear canals, respectively. **Tone-pip stimuli** had a duration of 100 ms, including 5 ms ramps. The stimulus level was 40 − 50 dB SPL. The range of stimulus frequencies was 1 − 9 kHz, with a typical step size 200 − 500 Hz, and 3 − 20 repetitions for each stimulus used. **Clicks** were presented at attenuation levels 55 − 0 dB, calibrated to correspond to stimulus levels 10 − 65 dB SPL, respectively (128 − 3300 repetitions at each single-unit recording location). Condensation clicks had a rectangular form, a duration of two samples (equivalent to 41.6 *µ*s) and an inter-stimulus-interval of 500 ms. **Spontaneous activity** was recorded for about 15 − 60 minutes for each unit.

#### Intracranial methods and recording protocol

Tungsten electrodes with impedances 2 − 20 MΩ were used (F.C. Haer, Bowdoin, ME). A grounded silver chloride pellet, placed under the animal’s skin around the incision, served as the reference electrode (WPI, Sarasota, FL). Electrode signals were amplified and band-pass filtered (10 − 10, 000 Hz) by a custom-built headstage and amplifier. Amplified electrode signals were passed to a threshold discriminator (SD1, TDT) and an analogue-to-digital converter (DD1, TDT) connected to a workstation via an optical interface (OI, TDT). In all experiments, voltage responses were recorded with a sampling frequency of 48, 077 Hz, and saved for off-line analysis.

For an intracranial recording, an electrode was advanced into the brainstem guided by stereotaxic coordinates, and units were characterized based on recorded extracellular spikes. Units were recorded on both sides of the brain. At each recording site, frequency responses were measured for tonal stimuli to each ear, and ITD tuning was measured with binaural tonal stimuli. Recordings confirmed that responses within nucleus magnocellularis (NM) were monaural, as expected. Single unit frequency response curves were recorded for the ipsilateral stimulus: for each recording location, an appropriate range of stimulus frequencies (within 1 − 9 kHz) was selected to record iso-level frequency response curves. Between single-unit recordings, the electrode was moved typically in steps of 100 *µ*m while searching for the next unit. For some units there were additional control recordings in which the recording from the same unit was continued while moving the intracranial electrode with steps of size 10 − 20 *µ*m.

#### EEG methods

An EEG signal was recorded simultaneously with all the intracranial recordings. Recordings were made using two platinum subdermal needle electrodes (Grass F-E2; West Warwick, RI) on the scalp. EEG signals were amplified using a WPI DAM-50 extracellular preamplifier, 0.1 − 10, 000 Hz (World Precision Instruments, Sarasota, FL). The EEG signal was further amplified (100x) using a custom built amplifier, and digitized (DDI, TDT). The voltage responses were recorded with a sampling frequency of 48, 077 Hz and saved for off-line analysis.

The active EEG electrode was always positioned in the dorsal midline, adjacent to the craniotomy, and the EEG reference electrode was positioned behind the ipsilateral ear. EEG electrodes could be slightly repositioned during the recording session to improve the signal.

### Intracranial recordings: Data analysis

In addition to custom Matlab scripts, we used the XdPhys script from M. Konishi’s lab and the supramagnetic wavelet-based ‘Wave-clus’ method for spike detection and clustering (Quian Quiroga et al., 2004), as provided as a Matlab script at https://github.com/csn-le/wave_clus.

#### Spike detection and clustering

We recorded from 151 intracranial locations within the NM cell body region (Fig. 1A) on which the spike detection and clustering was performed. Spikes were detected off-line, and all the data from a single intracranial recording location were combined. After spike detection and clustering, the spikes were categorized by their respective stimulus conditions (tone, click, spontaneous).

**Figure 1:**
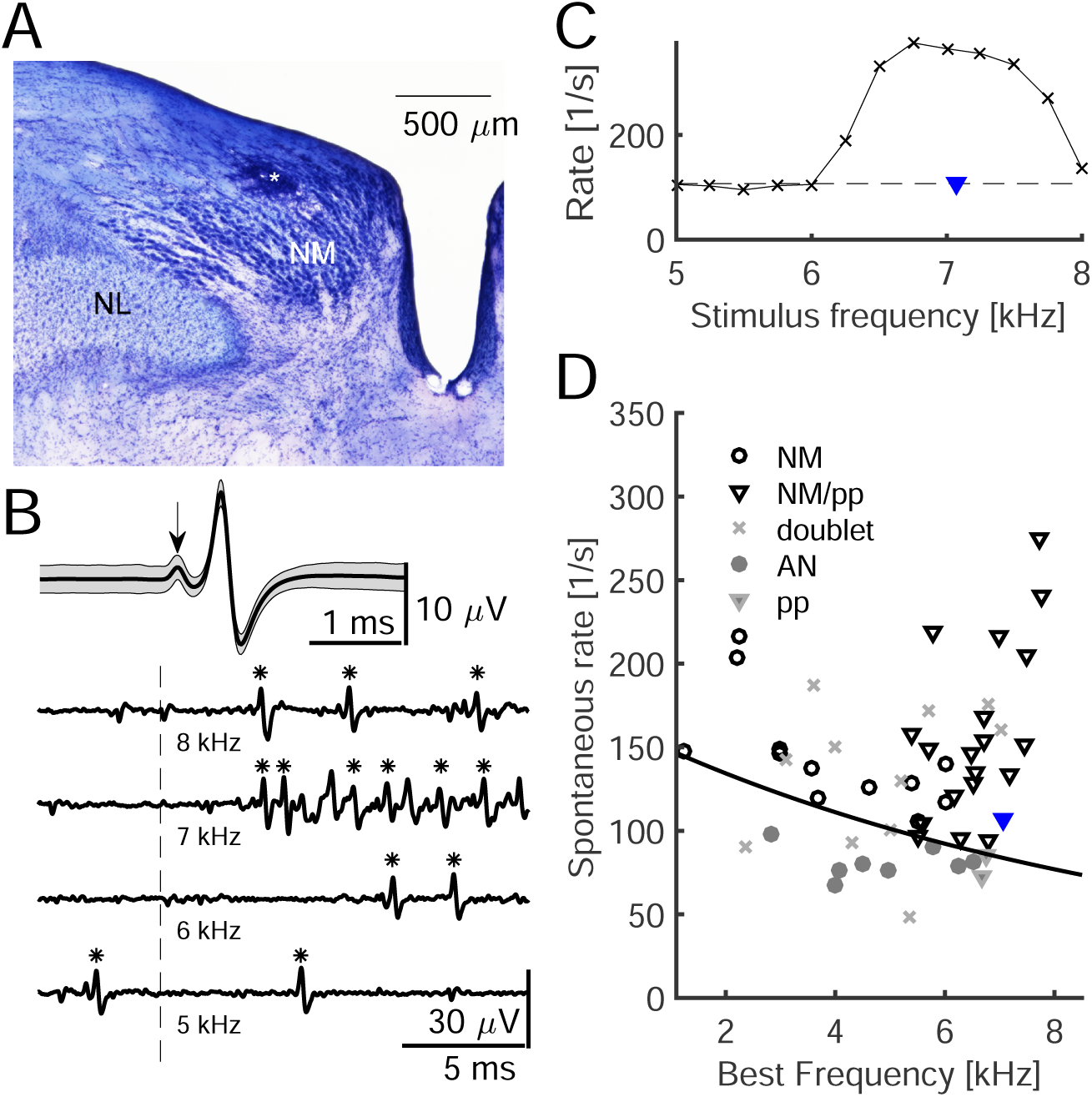
Recordings from NM cell body region. **A**: Exemplar recording location (lesion, *) in a Nissl-stained coronal slice through the auditory brainstem. The nucleus laminaris (NL) is both ventral and lateral to NM. **B**, Top: Average waveform of 22 641 spontaneous spikes (black line) ± SD (gray backgound); prepotential indicated by arrow. Bottom: Extracellular recordings from an NM neuron in response to tones at different frequencies (tone onsets indicated by vertical dashed line, detected spikes marked with *). **C**: Frequency-response tuning curve to pure tones at 50 dB SPL, with a maximum driven spike count rate of 376 spikes/s at 6750 Hz stimulus frequency. The best frequency (BF, marked with a blue triangle) of this unit was 7065 Hz. The dashed line indicates the spontaneous spike count rate 107 spikes/s. **D**: Spontaneous firing rates and BFs of all 53 units. Legend: *NM* : nucleus magnocellularis unit without a prepotential. *MN/pp*: nucleus magnocellularis unit with a prepotential. *AN* : auditory nerve fiber unit. *pp*: low-spontaneous rate unit with a prepotential. *doublet*: any unit with doublet-spiking. The NM/pp-unit shown in B and C is marked additionally with a blue triangle. Solid line: the decision boundary between NM and AN units (see Materials and Methods).

##### Spike detection

For spike detection, the default parameters of the *Wave-clus* method (Quian Quiroga et al., 2004) were modified as follows: The minimum threshold of spike detection (parameter ‘std_min’) was set manually for each unit depending on its spike size and noise level, and varied between 3.0 and 8.0 standard deviations (SD). Also the polarity of the spikes (‘detection’) was set manually for each unit upon visual inspection, because our set-up allowed spikes having either polarity. For the spike detection, the band-pass filter setting was 900−6, 000 Hz (‘detect_fmin’ and ‘detect_fmax’, respectively). The window length for spike shape was 1 ms before the spike peak and 1.5 ms thereafter, corresponding to ‘w_pre’ = 50 and ‘w_pre’ = 75 samples. The refractory time for the detection was set to 0 ms (‘ref_ms’), firstly because instantaneous firing rates in NM can be as high as 1, 500 spikes/s (Carr and Boudreau, 1993), and secondly because then we could detect units with spike-doublets. The ISI distribution of each unit was later scrutinized to exclude multi-units and doublet-units (see section ‘Prepotentials and doublets’).

##### Spike clustering

The spikes were clustered with the wavelet decomposition method within *Wave-clus* with 5 ‘scales’ in the wavelet decomposition and minimum of 10 inputs per cluster (‘min_input’). The radius of the clustering (‘template_sdnum’) was set to 4.5 SD, and the number of nearest neighbors (‘template_k’) was set to 10. Otherwise, both for detection and for clustering, the default parameters were used. After visual inspection of the resulting spike shape clusters, the clusters were merged if necessary (typically 2 − 3 clusters with an identical spike shape but variability during the onset or offset within the spike-window). Recording sites containing several units (with variable spike waveforms) were discarded from further analysis. In some recordings there was a small number of outliers (detected peaks not fitting any spike cluster) with always *N_out_ <* 0.75% of number of spikes in the main cluster(s); typically *N_out_* = 0 − 50. These outliers were excluded from the analysis.

##### Spike separation to stimulus conditions

**Tone-driven spikes**, obtained in response to 100 ms tones and with ≥ 15 dB SPL stimulus level, were included in the analysis when they occurred within 15 − 95 ms of the stimulus onset, thus excluding possible onset and offset effects. The **click responses** of the single-unit activity (*peri-stimulus time histogram, PSTH*) were calculated within 0 − 10 ms of the click stimulus onset. We considered **spontaneous spikes** to be any activity in trials in which there was no stimulus presented. Additionally, to collect as many spontaneous spikes as possible, we considered spikes to be spontaneous in two scenarios: Spikes occurring in stimulated trials (1) but later than 50 ms after the end of tonal or click stimuli, and (2) during stimuli that did not evoke an elevated sustained response, i.e. low-amplitude tones *<* 15 dB SPL at frequencies far off from the best frequency, excluding the first 20 ms after the stimulus onset.

##### Exclusion of recordings

We excluded units using three criteria: (1) Units with too few spontaneous spikes recorded (*<* 5000) because in this case we could not derive a significant spike-triggered average EEG (STA EEG, see Materials and Methods subsection, ‘EEG electrode recordings: Data analysis’). (2) Units for which the single-unit isolation was poor, i.e., the spike waveform SNR was *<* 8.6 dB. The SNR of the spike waveform was defined by the squared ratio of the spike peak amplitude and the standard deviation of the baseline. (3) Units for which the single-unit isolation broke down at the onset of the click-stimulus as confirmed by a visual inspection (see also ‘Click-evoked magnocellular activity’ below). After applying these exclusion criteria on the 151 units recorded within NM, 53 single units remained and were further analyzed.

#### Classification of magnocellular and auditory nerve units

Single units recorded within NM were classified to be either ‘AN fibers’ or ‘NM cell bodies /axons’; classification was based on best frequency (BF) and spontaneous firing rate, which were defined as follows:

##### BF

Iso-level response curves of the numbers of spikes defined the BF at a recording site as follows (Kuokkanen et al., 2010): a line at half height of a tuning curve was derived from its peak rate and the spontaneous rate. The midpoint of the line at half height yielded the BF. The best frequencies ranged from 1.25 to 7.75 kHz with mean ± SD: 5.60 ± 1.60 kHz. The tuning was calculated for the sustained activity in the window of 15 − 95 ms after tone onset, across all repetitions of the stimulus.

##### Spontaneous rate

Spontaneous rate was defined as the reciprocal of the mean spontaneous inter-spike-interval.

Auditory nerve and NM categories were based on the spontaneous firing rates and the characteristic frequencies (CFs) reported in Köppl (1997a), which provides the fits of CF vs spontaneous rate, *S*, for AN (*S*_AN_ [spikes/s] = 123.8 · exp(−0.129 · CF [kHz])) and NM units (*S*_NM_ [spikes/s] = 255.1 · exp(−0.0634 · CF [kHz])). We used the separating line of *f* · *S*_NM_ + (1 − *f*) · *S*_AN_ with *f* = 0.3, as this was the best line of separation between the AN / NM classes in Köppl (1997a).

#### Prepotentials and doublets

Recordings of magnocellular units can exhibit both prepotentials (Zhang and Trussell, 1994) and spike-doublets (Carr and Boudreau, 1993; Kuokkanen et al., 2018). For our analysis, a recording with a prepotential was interpreted as the intracranial electrode being located in the vicinity of an NM cell body and at least one large synapse from AN to this NM cell. In recordings from NM units, also spike-doublets can occur with very short inter-spike-intervals (ISIs 0.22 − 0.5 ms, Carr and Boudreau (1993), their Fig. 2). However, units with doublets pose a challenge both for spike sorting as well as for the estimation of the STA EEG, especially because the STA EEG waveforms are expected to temporally overlap for spike doublets.

**Figure 2:**
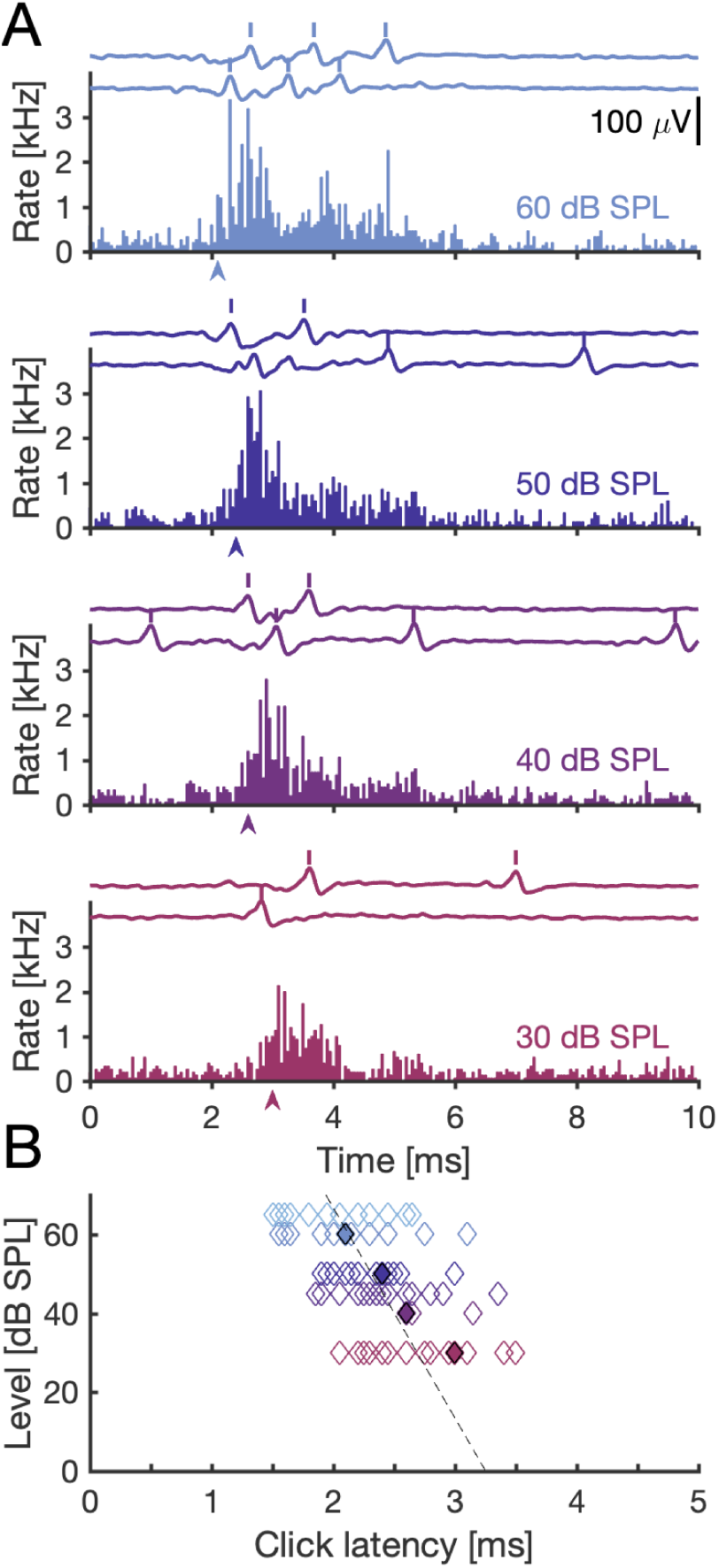
Click-response latency in NM is level- and BF-dependent. **A:** Peri-stimulus time histograms (PSTHs) obtained at four different levels of click stimuli from an extracellularly recorded single NM unit. Each PSTH is a summary of responses to many clicks. At the top of each panel, we show two voltage traces from example trials with spike times marked by vertical bars, indicating the low number of spikes in each single trial and the high variability across trials, as expected (Köppl, 1997a; Fontaine et al., 2015). Bin width: 50 *µ*s. The arrow-heads mark the click-response latency at each level. **B:** Click-response latency decreased with increasing stimulus level and with increasing BF. The examples in A are marked with filled diamonds. Dashed line: −19 ± 3 *µ*s/dB · level + 3.3 ± 0.2 ms (the GLM for the mean BF = 5.58 kHz). 32 NM units, with 1–4 stimulus levels each, resulting in *N* = 91 click-response latencies.

Upon visual inspection, 8 NM units were determined to include a large proportion of doublets and were excluded from further analysis.

#### Click-evoked magnocellular activity

The click-evoked responses of the single units (*peri-stimulus time histogram, PSTH*) were calculated within 0 − 10 ms after click onset.

The onset delay (or ‘click-response latency’) of the PSTH characterized the click-evoked responses in NM. We calculated the click-response latency using the same criterion as Köppl (1997a) — the first PSTH bin (with a 50 *µ*s bin size) after the stimulus presentation exceeding the largest spontaneous PSTH bin and being followed by a bin also fulfilling this criterion was defined as the click-response latency.

### EEG electrode recordings: Data analysis

In the following, we use the term ‘electroencephalography’ (EEG) when talking about a recording technique or signals acquired by it. These signals may be recorded during spontaneous or driven activity. We use the term ‘auditory brainstem response’ (ABR) to describe the evoked EEG signal in response to a specific kind of acoustic stimulus, namely clicks. We used click stimuli because this is the stimulus often used in human ABR for diagnostic purposes; furthermore, the ABR in response to clicks has a higher signal-to-noise ratio than in response to tonal stimuli.

#### Auditory brainstem response (ABR) recordings

We recorded click-evoked responses at the EEG electrode, i.e. the ABR, within either 0 − 10 ms or 0 − 15 ms after click onset. ABR waveforms were averaged across stimulus repetitions (127 − 500 trials) resulting in a ‘trial-averaged ABR’ for unchanged recording and stimulus conditions.

ABRs were quantified by the SNR, which was defined as the squared ratio of the peak amplitude of the trial-averaged ABR and the mean RMS of the baseline across ABR trials (5 ms window prior the click onset). The SNRs across the ABR waveforms ranged from −50 dB to +18 dB, with median of +5 dB. After visual inspection, we excluded ABR waveforms with the SNR *<* −13 dB (*<* 7% of ABR waveforms) as well as ABR waveforms not showing three peaks in the waveform (*<* 2% of the waveforms). The excluded responses were typically, but not always, recorded with a low stimulus level.

##### ABR wave quantification

We quantified the timing and amplitude of 3 positive waves in each waveform objectively as follows: We band-pass filtered (550 − 4, 000 Hz, Chebyshev type II filter of the order 8) the trial-averaged ABR response, and zero-mean-centered the waveform, to remove the low- and high-frequency noise present in some ABRs. We then used the Matlab algorithm findpeaks.m to find all peaks in this filtered ABR within 0 − 10 ms after stimulus onset. The algorithm returns the locations and heights of the peaks, and also the Matlab variables ‘width’ and ‘prominence’ (width at the half-maximum with respect to the baseline of the individual peak, and the height of the peak with respect to the same baseline). To identify the possible ABR peaks, we included all the maxima exceeding the threshold of 0.4 SD of the trial-averaged preamplifier-filtered ABR response (0 − 10 ms after stimulus onset). The threshold was chosen such that at least 2 ABR peaks were detected for all the waveforms. Of all the peaks crossing the threshold, we excluded the peaks with a ‘width’ narrower than 0.1 ms because typical ABR waves are much wider. If more than three peaks crossed the threshold, we used the three peaks with the highest ‘prominence’. If only two peaks were initially detected, we assumed that these would correspond to the peaks of the waves I and II because they typically were the largest peaks of the ABR, whereas the peak of the wave III was often small or even negative with respect to the baseline; thus, we included the largest peak within the period of 0.4 ms after the second found peak’s timing (starting point) to 3.0 ms after the first peak’s timing (end point). The starting point was selected to ensure that occasional small, local maxima within the wave II were not included, and the end point was selected because 3 ms was the typical duration of the ABR waveform from the wave I peak to the large negativity after wave III. Finally to ensure not to introduce jitter to the peak times because of the filtering, we applied these peak locations to the original preamplifier-filtered, trial-averaged ABR by finding the related maxima, allowing for a change of peak time of at most ±3 data points. In the end, this algorithm allowed us to quantify three peaks for all the selected ABR recordings.

The peak amplitudes of waves I to III were calculated from the preamplifier-filtered average traces, in comparison to the trial- and time-averaged baseline in the time window from the beginning of the recording (5 − 10 ms prior to click onset) to the time point 1 ms prior to wave I peak.

##### ABR averaging

The trial-averaged ABRs, as just defined, were obtained for different EEG electrode positions, intracranial recording sites, and click levels. After the ABR wave quantification, we averaged the detected peak amplitudes and their timing across trial-averaged ABRs for constant click levels as follows:

1. For the ABR waveform analysis tied to NM single units, all the trial-averaged ABRs recorded simultaneously with the respective single unit responses were used (1 − 11 trial-averaged ABRs with median of 1, in total 128 − 3300 trials, median: 300). For each NM unit the EEG electrode position was kept unchanged.
2. For the ABR waveform analysis unrelated to NM single units, we averaged peak amplitudes and their timings also across intracranial recording sites (1 − 14 trial-averaged ABRs with median of 1, in total 128 − 4200 trials, median 999). In some days the EEG electrode was re-positioned during the experiment. Here, we restricted the ABR waveform analysis to the EEG electrode position with the highest signal-to-noise ratio (SNR), resulting in *N* = 24 EEG electrode positions.

##### ABR inter-peak-intervals

The inter-peak-intervals of peaks 1 − 2, 1 − 3, and 2 − 3 were calculated based on the delays of peak timings in trial-averaged ABR waveforms and thereafter averaged as described above.

#### Spike-triggered average EEG

EEG traces recorded in the absence of acoustic stimuli were band-pass filtered (800 − 3000 Hz, Chebyshev type II filter of the order 6). Compared to the ABR recordings, a narrower filter was chosen to further reduce noise. The spike-triggered average EEG (STA EEG) was calculated for each NM single unit separately. The STA EEG was derived from 8-ms time windows (*N_t_* = 402 data points) of the EEG recording centered at spike times of single units. We used only spontaneous spikes for the STA EEG.

We define the STA EEG mathematically as the average signal at the EEG-electrode, *C*(*τ*), around isolated spikes of a neuron *j* at times *t^j^*(maximum voltage) where *i* = 1, 2*, …, n* is the spike number and *r*(*t*) is the simultaneously recorded EEG:

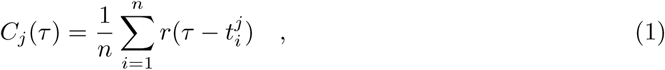

using the interval of −4 ms ≤ *τ* ≤ 4 ms around each spike for the analysis.

We excluded from further analysis units by two criteria as follows: 1) To ensure that the EEG signal was high enough for the calculation of the STA EEG and for the calculation of the NM single-cell contribution to the ABR, the SNR of the ABR waveform at the highest click levels was required to be ≥ 1 dB, leading to exclusion of two NM single units. The SNR of the STA EEG waveform was defined by the squared ratio of the spike peak amplitude and the standard deviation of the baseline.

2) To ensure that there was no cross-talk between the intracranial and EEG electrodes, we excluded the 7 units (out of 31) with an SNR *>* −18 dB of the STA EEG (range: −79 to +6 dB). This led to an SNR of the STA EEG of *>* −15 dB. In these units, the average spike waveform of the intracranial electrode and the waveform of the STA EEG were practically identical. After exclusion of 7 NM single units with putative crosstalk in the STA EEG, there were 24 NM single units included in further analysis.

##### STA EEG waveform significance

The significance of the STA EEG waveform was judged by using two bootstrapping methods. Firstly, the significance of the waveform was estimated with the SNR-based bootstrapping method by Parks et al. (2016). We note that this SNR has a different definition than the one in the previous paragraph: Here the number of samples was the number of spontaneous spikes, and the SNR distribution was based on 9999 bootstrap samples. The post-window width, for which the *signal* is calculated, was ±0.25 ms around the spike time, corresponding to a post-window width of 0.5 ms. The pre-window width, from which the respective *noise* level is calculated, was set to 1.75 ms, (from 4 ms to 2.25 ms before the spike time). The 10-percentile lower bound threshold was set to 0 dB based on our SNR distributions. We chose a rather short post-window width to avoid being overly selective about the units left for the prediction of the ABR contributions (see below).

After establishing which of the STA-waveforms as such were significant, the time points (from −1.4 to 1.0 ms with respect to the spike time) at which each was significant were identified as by Teleńczuk et al. (2015), with the 2-sample bootstrapping method with the confidence interval of 99% of the SE. There was no correction for multiple comparisons.

### Control experiment

We conducted control experiments to confirm that electrical cross-talk between the EEG and intracranial electrodes in general did not affect our results. The idea behind these control experiments is as follows: when the intracranial electrode is moved the intracranial spike waveform changes. If there is cross-talk between the intracranial and the EEG electrodes, the STA EEG waveform should change as well. In contrast, if there is no cross-talk, the STA EEG should be independent of the intracranial spike waveform.

We thus moved in an exemplary control experiment the intracranial electrode in ten steps of 10 − 20 *µ*m over a total distance of 120 *µ*m in the vicinity of an NM cell body. At the initial recording depth, the spike amplitude was 24.21 ± 0.02 *µ*V (mean ± SE; the spike waveform and the related STA EEG from the initial recording depth is shown in the later Figure 4A). Moving the intracranial electrode deeper into the tissue changed the peak amplitude of the spike. After the first 10 *µ*m-step, the spike amplitude peaked at 26.19 ± 0.04 *µ*V and then decreased monotonically to 14.74 ± 0.03 *µ*V (120 *µ*m away from the first recording position). The amplitude of the prepotential behaved similarly, starting at 4.96 ± 0.02 *µ*V, peaking after 10 *µ*m at 5.39 ± 0.04 *µ*V, and then decreasing monotonically to 2.77 ± 0.03 *µ*V. The relative delay between the prepotential and the spike peak monotonically increased from 460 *µ*s to 660 *µ*s with depth. The spike amplitude and the prepotential amplitude were significantly correlated with the recording depth and with each other: the Pearson correlation coefficient between the spike amplitude and prepotential amplitude was 0.98 (*p* = 4.0 · 10^−7^), the correlation between spike amplitude and depth was −0.72 (*p* = 0.019), and the correlation between prepotential amplitude and depth was −0.77 (*p* = 0.0097).

By contrast, the STA EEG waveform did not change significantly when the intracranial electrode was moved. The change was evaluated as follows: There was always a significant positive peak at −190 *µ*s and always a significant negative peak at 130 *µ*s delay (*p <* 0.05 for each intracranial depth, SD bootstrapping method). Interestingly, the peak amplitudes were independent of the intracranial depths: the Pearson correlation coefficient between the STA EEG amplitude and recording depth was 0.49 (*p* = 0.15) for the positive peak at −190 *µ*s delay and −0.09 (*p* = 0.80) for the negative peak at 130 *µ*s delay.

In summary, the control experiment provides evidence against cross-talk between the intracranial and the EEG electrodes in general, and thus supports the absence of contamination between the intracranial electrode and the EEG electrode.

### Prediction of the single-unit contribution to the ABR

To predict the single-unit contribution to the ABR, we used the recordings from 24 NM units (in 11 owls). From the single-unit recordings obtained for click stimuli, we obtained the trial-averaged peri-stimulus time histograms (PSTHs), which we mathematically describe by the function PSTH*_j_*(*s*) for neuron *j* = 1*, …,* 24 for 0 ≤ *s* ≤ *T_j_* with click at time *s* = 0 and recording duration *T_j_* ∈ {10, 15} ms after the click onset). From the EEG recordings, we had obtained the ABR waveforms. And from the combined intracranial and EEG recordings during spontaneous activity, we had derived the STA EEG, i.e. *C_j_*(*τ*) for neuron *j* and *τ* ∈ [−4, 4] ms, in Equation (1).

To calculated a single NM cell’s contribution to the ABR in one trial, we sum its STA EEG waveforms at the given spike times for this trial. Summing the single-trial contributions across all stimulus trials and normalizing by the number of trials gives us the average contribution of the unit in any trial, which we denote as ABR*_j_*(*t*) for neuron *j*. We can then reorganize this sum to a convolution: To predict a single NM cell’s average contribution to the ABR, we convolved the PSTH of that neuron with its STA EEG:

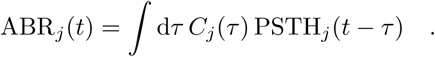

#### Statistical analysis

All analysis was performed with a custom-written MATLAB code. To estimate the statistical significance of the data, we used the Pearson correlation coefficient and its p-value, N-way analysis of variance (ANOVA), generalized linear models with respective F-statistics, Student’s 2-population t-test, and custom bootstrapping methods as explained across Materials and Methods. When correction of multiple testing was done, we used the Šidák correction (Abdi, 2007).

#### Data availability

All the data and codes used to produce the figures in this study are available from the corresponding author upon request.

## Results

The aim of this study was to quantify the contribution of the auditory brainstem nucleus magnocellularis (NM) to the auditory brainstem response (ABR). To this end, we determined the contribution of single neurons to the ABR by recording action potentials in NM units simultaneously with the EEG from the scalp. These simultaneous recordings allowed us to estimate the spike-triggered averages (STAs) of NM neurons at the EEG electrode (i.e., the unitary responses). Having measured the click-evoked spike times of the same NM neurons, we could then estimate the neurons’ contribution to the click-evoked EEG response, i.e., the ABR.

### Classification of single units

To link single cell activity to their contributions to the EEG signal, we analyzed extracellular recordings from 53 single units in 12 owls, obtained within the NM cell body region (Fig. 1A,B). This region also contains auditory nerve (AN) fibers that descend into NM, and NM efferent axons. Thus, AN fibers, NM cell bodies, and NM axons could, in principle, have been recorded at any of the depths used. We classified these units based on their best frequency (BF) and spontaneous firing rate (Fig. 1C,D), since AN units typically have lower spontaneous rates (for each BF) than NM units (Köppl, 1997a). Based on these earlier results, 13 units were putatively classified as AN and 40 units were classified as NM.

We also used the presence and absence of prepotentials (example in Fig. 1B, top) to differentiate between NM cell bodies and AN fibers (see Materials and Methods). Prepotentials have been observed in avian endbulb synapses between AN and NM (Zhang and Trussell, 1994). In the mammalian auditory system, prepotentials originate from the large endbulb of Held synapse between the AN and the anterior ventral cochlear nucleus (AVCN) and from the calyx synapse in the medial nucleus of the trapezoid body (MNTB) (see Discussion, e.g. Pfeiffer, 1966; Kopp-Scheinpflug et al., 2003; Englitz et al., 2009). We concluded that single-units with a prepotential originated, with a high probability, from the vicinity of NM cell bodies (see Table 1 for their properties). Most units with a prepotential (19 out of 21, black downward open triangles in Fig. 1D) aligned with our classification as NM that was based on BF and spontaneous rate. The two units with a low spontaneous rate but showing a prepotential (gray filled downward triangles) were classified as ambiguous (see Materials and Methods).

**Table 1:** Descriptive statistics of the NM population. Abbreviations: SNR: signal-to-noise ratio. ISI: inter-spike interval.

| Variable | mean ± SE | range | $N$ |
| --- | --- | --- | --- |
| Number of spontaneous spikes | 43 700 ± 1 100 | [10 558, 140 141] | 32 |
| Spontaneous rate (1/mean ISI) | 151 ± 2 spikes/s | [94, 275] spikes/s | 32 |
| Amplitude of spontaneous spikes | 13.4 ± 0.3 μV | [0.7, 28.4] μV | 32 |
| SNR of spontaneous spikes | 13.86 ± 0.07 dB | [8.91, 18.24] dB | 32 |
| Best frequency (BF) | 5580 ± 60 Hz | [1250, 7750] Hz | 32 |
| Prepotential amplitude | 2.20 ± 0.07 μV | [0.11, 4.96] μV | 19 |
| Prepotential amplitude, % of spike | 13.1 ± 0.3 % | [5.4, 21.2] % | 19 |
| Prepotential SD % of spike SD | 107.6 ± 1.2 % | [82.5, 174.9] % | 19 |
| Prepotential delay wrt. spike | 509 ± 6 μs | [340, 820] μs | 19 |
| Mode of ISI distribution | 900 ± 30 μs | [400, 2100] μs | 19 |

The stringent classification criteria used so far resulted in the identification of 40 units as originating from NM neurons. Among them, eight units were excluded because of a high proportion of spike doublets (gray crosses; see also Materials and Methods) because it is challenging to determine STAs for such units. Thus, 32 NM units from 12 owls were used in later analyses (black circles and black downward triangles in Fig. 1D; see also Table 1 for properties of these units).

### Click-evoked activity in NM

To evaluate the contribution of single units to the ABR, typically evoked by a click stimulus, we recorded peri-stimulus time histograms (PSTHs) of NM units in response to clicks. We recorded responses to a range of click levels for each unit (10 − 65 dB SPL, examples in Fig. 2A). To characterize the click-evoked single-unit responses from NM units, we described their single-unit PSTHs by click-response latency (arrowheads in Fig. 2A; for a definition of click-response latency, see Materials and Methods: ‘Click-evoked magnocellular activity’). This click onset timing could only be identified for clicks at ≥ 30 dB SPL.

At the population level, the NM units’ click-response latency decreased with increasing level (Fig. 2B) and with increasing BF. A generalized linear model (GLM) describing the click-response latency as a first-order polynomial of level and BF, with offset and linear link function, showed a significant dependence of click-response latency on both level and BF: −19 ± 3 *µ*s/dB · level − 90 ± 20 *µ*s/kHz· BF +3.8 ± 0.2 ms, with *p*(level) = 1 · 10^−8^ and *p*(BF) = 2 · 10^−5^ (F-statistics: vs. constant model: *F*_3,88_ = 34.4, *p* = 9 · 10^−12^; normally distributed residuals, no interaction term between BF and level: *p* = 0.61). If we neglect the dependence on BF, the level dependence of click-response latency had a slope of −19 ± 3 *µ*s/dB (Fig. 2B, dashed line). Köppl (1997c) reported similar values, showing delay-to-level slopes for the tone-elicited delays in 3 NM cells, with slopes ranging from −24 *µ*s/dB to −16 *µ*s/dB (fitted from their Fig. 9).

### ABR timing: Delays originate in the inner ear

In order to relate the activity of the single units to the EEG, we first measured and quantified the properties of the EEG on its own. We recorded ABRs in response to click stimuli whose sound levels varied from 10 to 65 dB SPL.

ABRs typically contained three positive-going waves within the first 8 ms following the click presentation (Palanca-Castán et al., 2016), and the latencies of the peaks of the three waves increased with decreasing stimulus level (examples in Fig. 3A). To quantify the dependence of the latencies of the peaks on the stimulus level, we analyzed the shift of the three waves as well as their inter-peak-intervals in 27 ABR recordings in 13 owls. The latency of the peak of each wave was indeed level-dependent (all Pearson correlation coefficients *<* −0.83 with p-values *<* 10^−20^) across the recordings, and their slopes (Fig. 3B) were not significantly different (GLM with mean-shifted intercepts, all *p* = 1, GLM: *F*_4,225_ = 231, *p* = 7 · 10^−68^). The level-dependent slope across all peaks was −23.1 ± 0.9 *µ*s/dB, with intercept 2.82 ± 0.05 ms for the first peak, 3.54 ± 0.04 ms for the second peak, and 4.49 ± 0.04 ms for the third peak (GLM: *F*_4,221_ = 1230, *p* = 2 · 10^−137^, mean ± SE).

**Figure 3:**
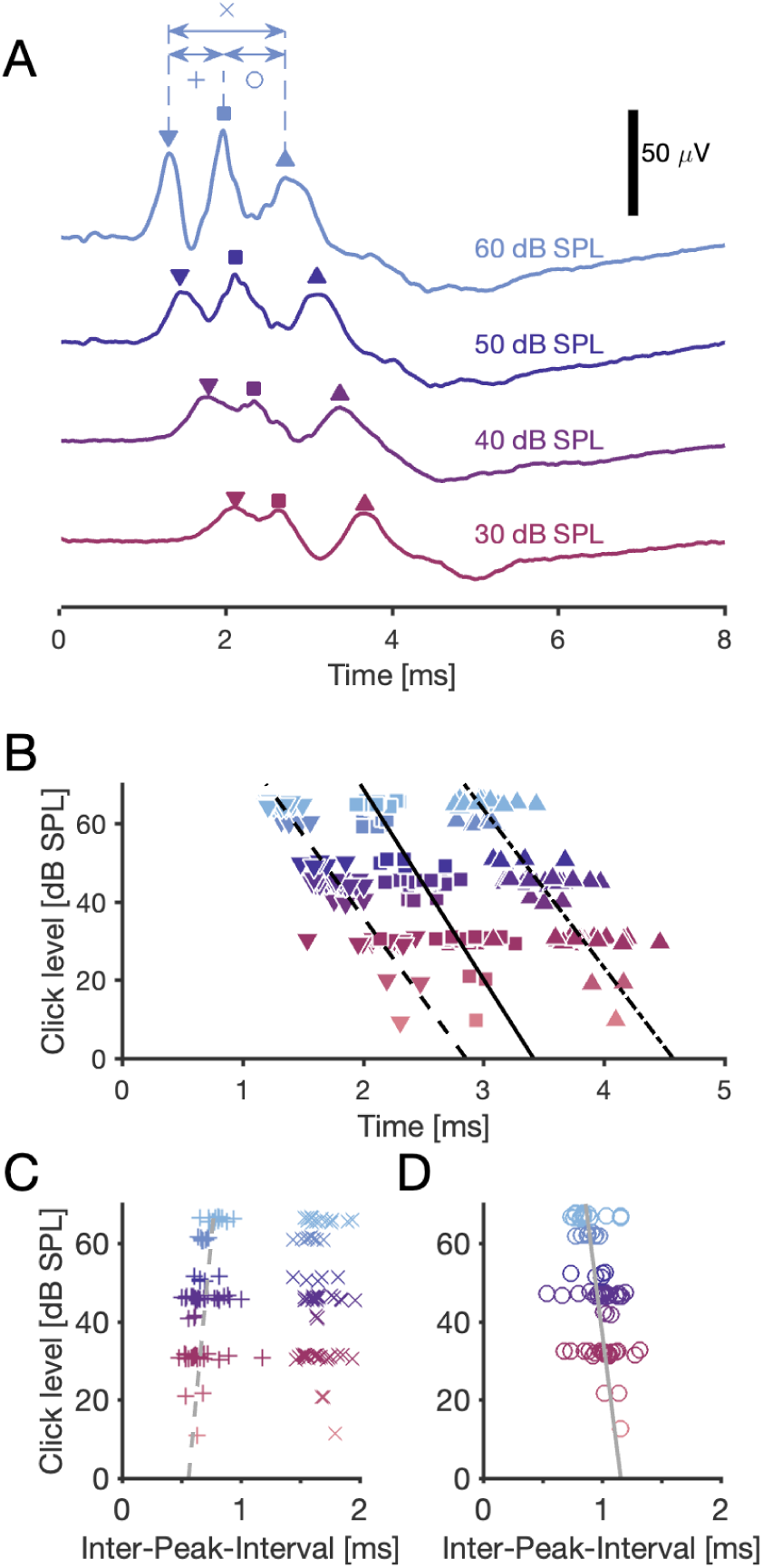
Delay of ABR waves depends on sound level. **A**: Examples of an ABR, recorded in response to four different levels of a click with onset at 0 ms. Each curve shows three main peaks (marked with symbols ‘∇’ for wave I, ‘□’ for wave II, and ‘△’ for wave III).e inter-peak-intervals are marked with symbols ‘x’, ‘+’, and ‘o’. **B**: ABR waves’ peak timing depended significantly on the stimulus level. Linear least-square fits (lines): Wave I peak: −24 *µ*s/dB · level + 2.853 ms. Wave II peak: −21 *µ*s/dB · level + 3.414 ms. Wave III peak: −25 *µ*s/dB · level + 4.573 ms. All groups: Pearson correlation coefficients < −0.84 with p-values < 10*^−^*^20^, *N* = 75 for each wave. The markers are jittered within 1 dB to reduce overlap. **C**: The inter-peak-interval between peaks 1 and 2 depended on the stimulus level as 3.1 *µ*s/dB · level + 0.561 ms (linear least-square fit), with Pearson correlation coefficient of 0.35 (*p* = 0.0022). The average inter-peak-interval (± SE) between peaks 1 and 3 it was 1.67 ± 0.02 ms with no significant correlation with level (Pearson CC: −0.11; *p* = 0.34). **D**: The inter-peak-interval between peaks 2 and 3 depended on the stimulus level as −4 *µ*s/dB · level + 1.159 ms (linear least-square fit), with Pearson correlation coefficient of −0.41 (*p* = 0.00034, *N* = 75). B–D: 24 ABR recordings, with 1–4 stimulus levels each, resulting in *N* = 75 delays and inter-peak-intervals per group.

The level-dependent fits for the click-response latency in the NM population (dashed line in Fig. 2B) and the ABR wave II peak delay (solid line in Fig. 3B) were equal within their error margins. We performed an N-way analysis of variance (ANOVA) based on the hypothesis that both groups (ABR wave II peak delay: *N* = 75 and click-response latency: *N* = 91) originated from the same level-dependent regression model. The group identity had no significant effect on the fit (*F*_1,156_ = 1.9, *p* = 0.18), whereas the level did (*F*_7,156_ = 19.2, *p* = 3 · 10^−18^), indicating that there was no significant difference between the delays of the ABR wave II peak and the NM cells’ click-response latency.

By contrast, the inter-peak-intervals (Fig. 3C,D) showed a much weaker level dependence. The inter-peak interval between peaks 1 and 3 (IPI_13_) showed no significant level dependence (Pearson correlation coefficient for IPI_13_: −0.0011 with *p*_1,3_ = 0.36, *N* = 75 in each IPI group), with mean (±SE) IPI_13_ = 1.67 ± 0.02 ms. The level dependency of IPI_12_ = 3.1 *µ*s/dB · level + 0.561 ms and of IPI_23_ = −4 *µ*s/dB +1.116 ms (linear least-square fits) were nevertheless significant (Pearson correlation coefficients of IPI_12_: 0.35, *p* = 0.0022 and of IPI_23_: −0.31, *p* = 0.0071).

Our results so far have implications for the origin(s) of the level dependence of delays in the auditory pathway. ABR wave I is assumed to reflect auditory nerve activity (Corwin et al., 1982; Melcher and Kiang, 1996). Consistent with this hypothesis, the strong overall level dependence of ABR latency in our data set was mainly defined by the response of the cochlea, which is level dependent. Furthermore, the much weaker dependence of inter-peak-intervals suggests that delays between brainstem nuclei are mainly caused by fixed structural delays, such as synaptic delays and axonal conduction delays, which are basically level independent.

Finally, we also quantified how the peak amplitude of the ABR wave II was modulated by stimulus level. The ABR wave II peak amplitude correlated in the population strongly with the level (Pearson correlation coefficient: 0.65, *p* = 4 · 10^−10^, *N* = 75) with the slope of 0.47 *µ*V/dB and intercept of −9.3 dB (linear least square fit).

### Spontaneous spikes of individual NM neurons were detectable in the EEG signal

To connect the action potentials of single NM cells to the macroscopic EEG, we analyzed the average EEG around the times of spikes. The average contribution of a spike from a single unit is referred to as spike-triggered average (STA) EEG. For this analysis we only used spontaneous spikes in order to avoid stimulus-induced correlations among neurons, which would distort the computed STA EEG. This was possible because the NM units have high spontaneous firing rates (Köppl, 1997a).

Eight NM units (of *N* = 32, Figs. 1 and 2) were excluded from the STA analysis because their respective EEG recordings failed the stringent inclusion criteria for the EEG; these criteria included both suspected crosstalk between the electrodes and weak ABR responses (see Materials and Methods).

Figure 4A,B shows two examples of NM units and their corresponding STA EEG. Two thirds of the analyzed NM neurons (16 out of 24) contributed a statistically significant STA EEG waveform (Fig. 4C) according to the SNR-method by Parks et al. (2016) with an SNR lower bound of 0 dB (see Materials and Methods). Averaging over thousands of spontaneous spike times per unit revealed significant waveforms (see text and asterisks next to the waveforms in Fig 4C). Across the population of 16 significant units, there was a large spread both in the amplitudes of the STA EEG peaks and their timing (Fig 4C, D). The peak amplitude of the STA EEG ranged from 25 to 267 nV (mean ± SE: 76 ± 4 nV, see Table 2). The noise in the EEG signal was typically about 3 orders of magnitudes larger than the peak amplitudes of the STA EEG, corresponding to a very low SNR of −42 ± 1 dB (mean ± SE, see Table 2).

**Figure 4:**
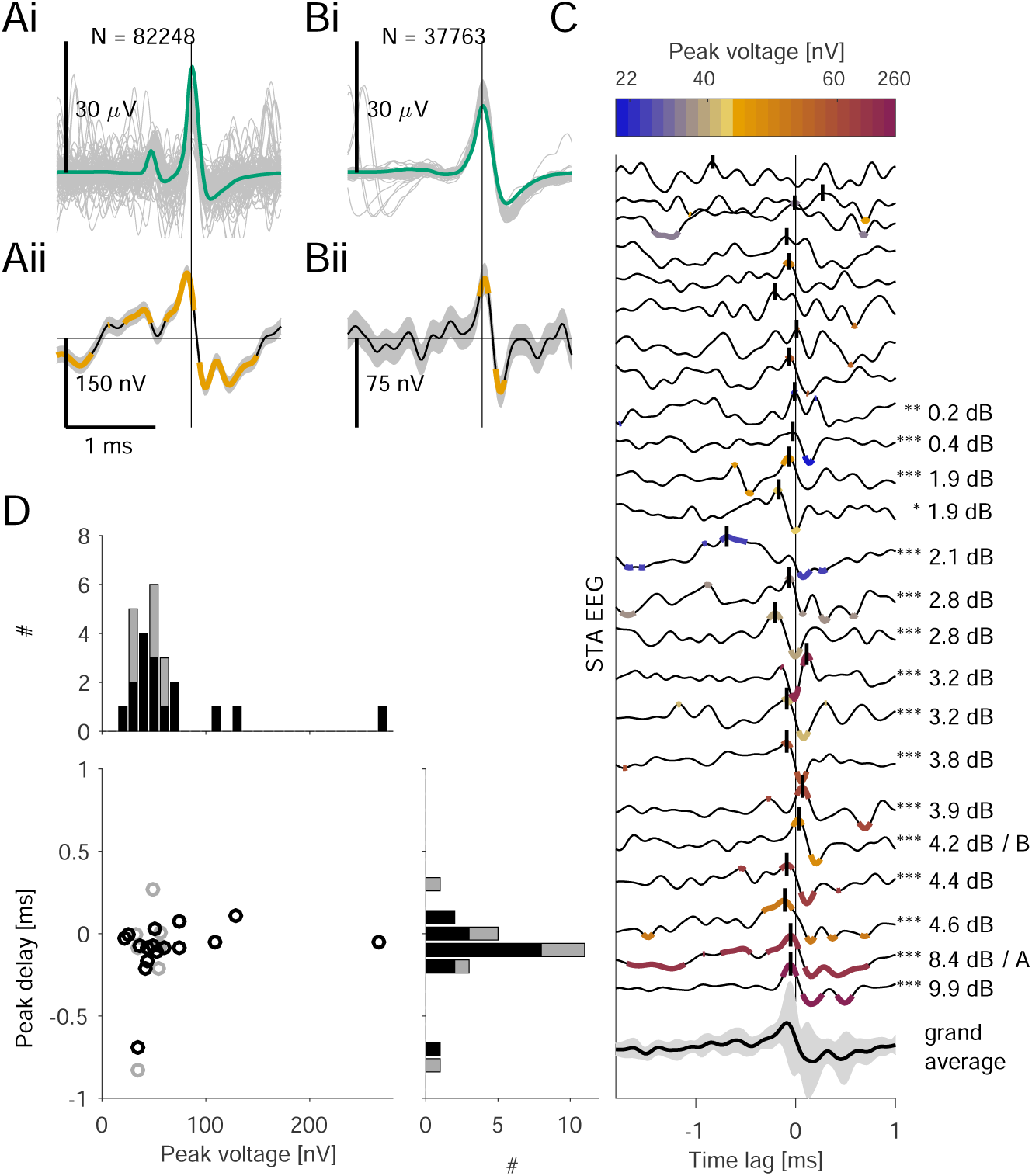
Magnocellular single cell spikes make a detectable contribution at the scalp electrode. **Ai:** Average spike waveform of 84 248 spontaneous spikes of an NM cell (green), recorded extracellularly, and a random selection of 100 spike waveforms thereof (gray). **Aii:** Average waveform at the EEG electrode (STA EEG, black) and SE (shaded), with EEG waveforms aligned to the peaks of the spikes of the NM cell in Ai (thin vertical black line). The parts of the STA EEG marked in orange have a significance level *p* < 0.01, and black portions are non-significant. **B:** Average spike waveform and STA EEG from a different NM unit. **C:** 24 STA EEGs, sorted by the significance of their peaks (vertical black bars) within ±1.0 ms with respect to the spikes of the respective NM units. Significant curves (SNR_LB_ ≥ 0 dB) are highlighted by black numbers of the corresponding values of the SNR_LB_ (*N* = 16); non-significant curves do not have values (*N* = 8). Asterisks indicate the maximum bootstrapped significance of the SDs of curves (*: *p* < 0.05, **: *p* < 0.01, ***: *p* < 0.001, see Materials and Methods), and significant parts of the waveforms are colored according to the colorbar at the top. Not significant parts are black. The grand average STA EEG (± SD in gray) of the significant curves is shown at the bottom, with the peak amplitude 40 ± 60 nV at −90 *µ*s. **D:** Peak delays (STA EEG wrt. the spike waveform) and maximum STA EEG amplitudes (peak voltages) were not correlated (Pearson CC: 0.20, *p* = 0.35, *N* = 24). Significant data points (SNR_LB_ ≥ 0 dB) are black (*N* = 16), and the non-significant ones are gray (*N* = 8). There was no difference between the two groups neither in the number of spikes, in the peak voltages, in the peak delays nor in the SNR of the STA EEG (see Table 2). Histogram on the top: distribution of the STA EEG peak voltages. Histogram on the right-hand side: distribution of the STA EEG peak delays. Population statistics: see Table 2.

**Table 2:**
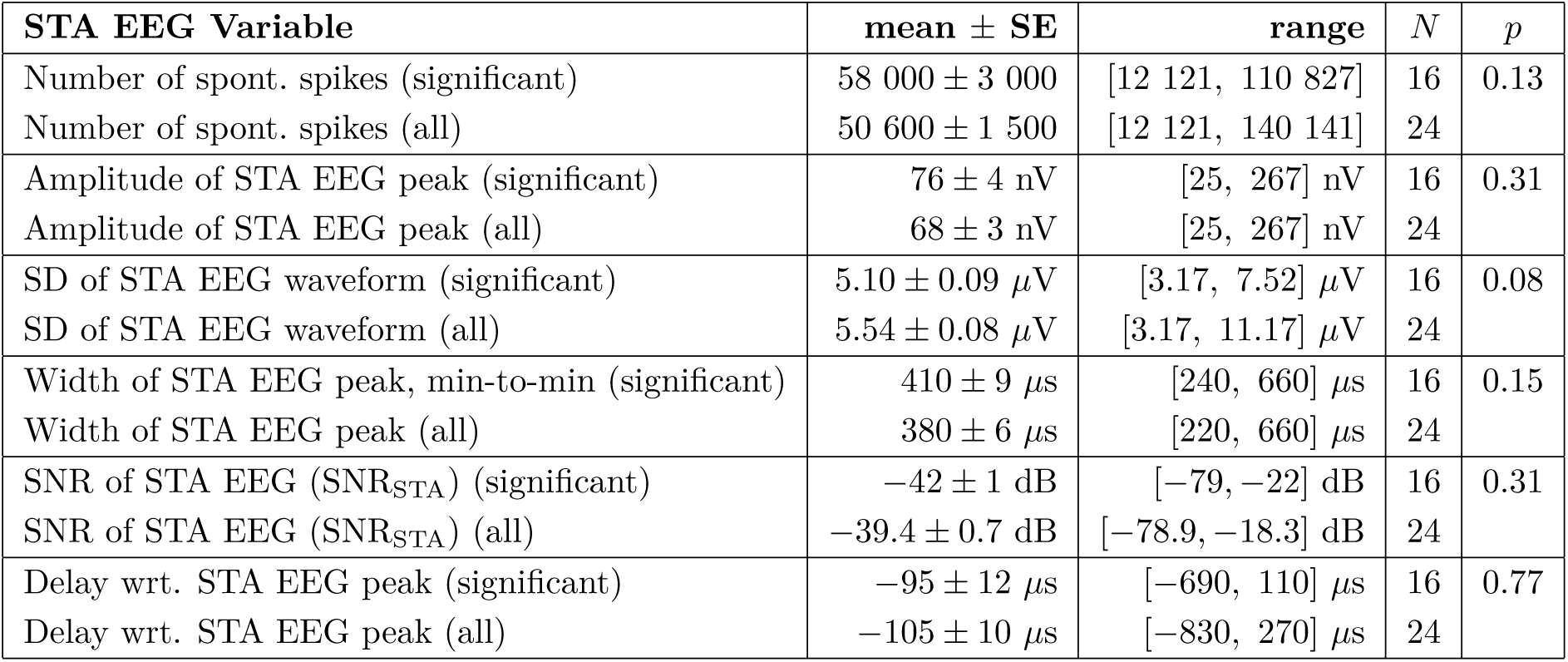
Spike triggered average EEG amplitudes and delays of NM units. The *p*-values refer to Student’s 2-population t-test between the STA EEG populations of significant (*N* = 16) and non-significant (*N* = 8) waveforms.

| STA EEG Variable | mean ± SE | range | $N$ | $p$ |
| --- | --- | --- | --- | --- |
| Number of spont. spikes (significant) | $58\,000 \pm 3\,000$ | [12 121, 110 827] | 16 | 0.13 |
| Number of spont. spikes (all) | $50\,600 \pm 1\,500$ | [12 121, 140 141] | 24 | |
| Amplitude of STA EEG peak (significant) | $76 \pm 4$ nV | [25, 267] nV | 16 | 0.31 |
| Amplitude of STA EEG peak (all) | $68 \pm 3$ nV | [25, 267] nV | 24 | |
| SD of STA EEG waveform (significant) | $5.10 \pm 0.09$ μV | [3.17, 7.52] μV | 16 | 0.08 |
| SD of STA EEG waveform (all) | $5.54 \pm 0.08$ μV | [3.17, 11.17] μV | 24 | |
| Width of STA EEG peak, min-to-min (significant) | $410 \pm 9$ μs | [240, 660] μs | 16 | 0.15 |
| Width of STA EEG peak (all) | $380 \pm 6$ μs | [220, 660] μs | 24 | |
| SNR of STA EEG (SNR<sub>STA</sub>) (significant) | $-42 \pm 1$ dB | [-79, -22] dB | 16 | 0.31 |
| SNR of STA EEG (SNR<sub>STA</sub>) (all) | $-39.4 \pm 0.7$ dB | [-78.9, -18.3] dB | 24 | |
| Delay wrt. STA EEG peak (significant) | $-95 \pm 12$ μs | [-690, 110] μs | 16 | 0.77 |
| Delay wrt. STA EEG peak (all) | $-105 \pm 10$ μs | [-830, 270] μs | 24 | |

Most of the STA EEG maxima occurred slightly prior to the maximum of the extracellular spike waveform, with a mean (± SE) delay of −95 ± 12 *µ*s (*N* = 16; see Table 2 and Fig. 4C, D). The STA EEG peak being close to the spike maximum is consistent with the assumption that we typically recorded intracranially close to the cell bodies and that the (far-field) dipoles originating from these neurons would have a similar but not necessarily equal peak time at the scalp.

### Predicted NM contribution matches the peak latency of the ABR wave II

To establish a direct connection between click-evoked NM single-cell activity (Fig. 2) and the ABR (i.e., click-evoked EEG response, Fig. 3), we recorded them simultaneously and used the STA EEG (Fig. 4) to predict the single-cell contribution to the ABR (Fig. 5). We did not attempt to explain the full ABR waveform, which is assumed to originate from all excitatory cells in the auditory brainstem (Achor and Starr, 1980; Corwin et al., 1982).

**Figure 5:**
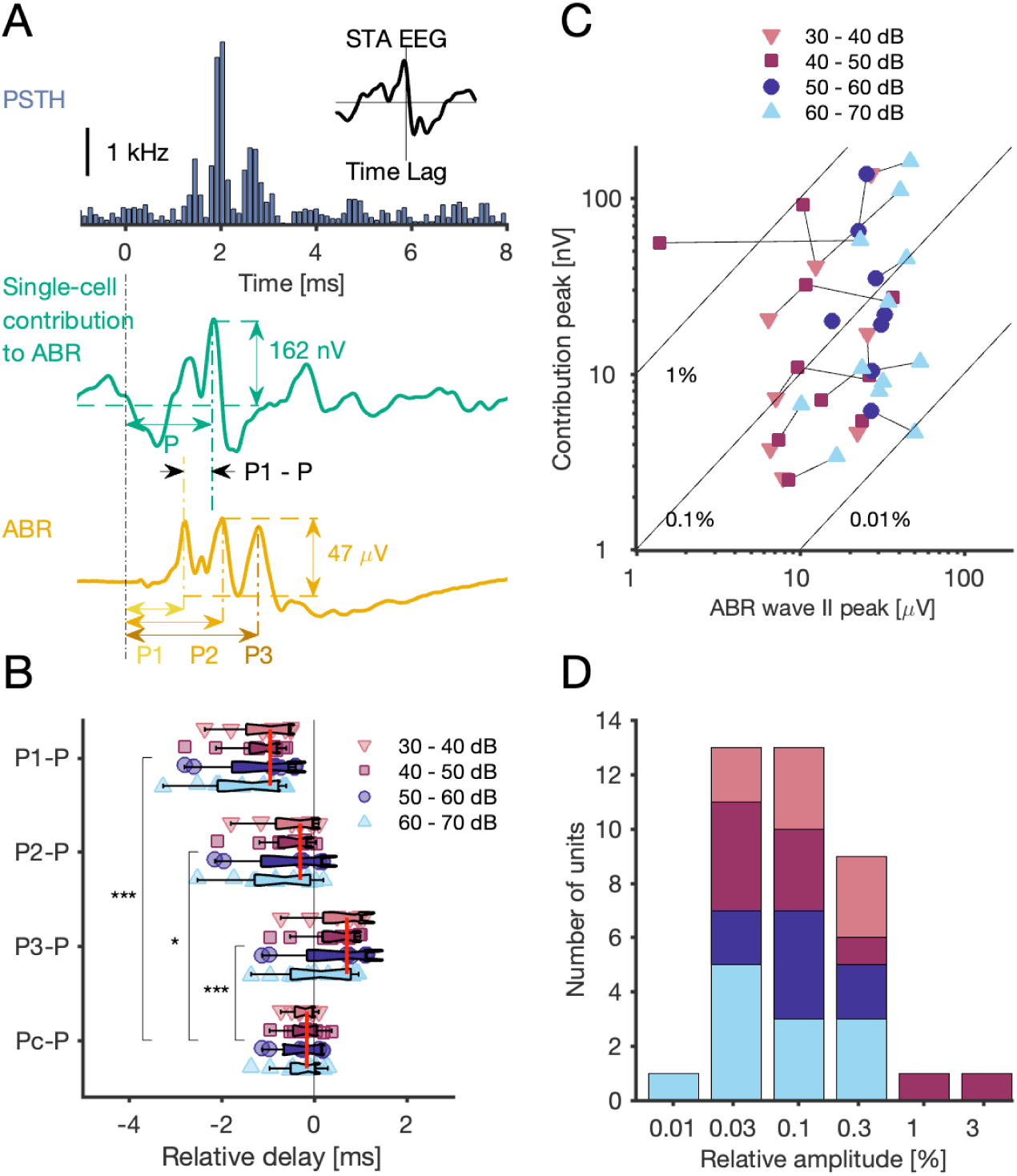
Predicted NM single-cell contribution aligns best with peak of ABR wave II. **A**, Top: PSTH (gray bars) in response to a click stimulus at 0 ms. Inset: STA EEG of the spontaneous spikes (*N* = 84 248; see Fig. 4Aii). Middle: Predicted single-unit contribution to the ABR (green), calculated as the convolution of the STA EEG with the PSTH (both shown above); peak amplitude of prediction: 162 nV (wrt. average level at click onset ±1 ms). Delay of peak indicated by ‘P’. Bottom: ABR (yellow) in response to the click stimulus; peak-to-peak amplitude of ABR wave II: 47 *µ*V (wrt. lowest neighboring minimum). Delay of peak indicated by ‘P2’. All parts of this panel share the same time scale, and the click onset is marked with a vertical line at 0 ms. **B–D**: Population data from 38 EEG recordings (at variable click levels) and from 16 NM cells. Plots share the same color schema with respect to stimulus levels (see legend in C). **B:** Boxplots and data points of the relative delays wrt. each ABR peak and for each level group. The relative delay is the difference between the delay *P* of the predicted single-cell ABR contribution peak and one of the delays (P1, P2, or P3) of a peak of ABR waves I through III; we also show the relative delay of the predicted peak and the closest ABR wave’s peak (*P_c_* − *P*; *: *p*= 0.011, ***: *p* < 0.0001, 2-population t-tests). The vertical red lines mark the medians of each relative delay across levels. **C:** Amplitude of predicted contribution peak vs. amplitude of ABR wave II. Short lines connect data points obtained from the same NM cell but at different click levels. Long diagonal lines indicate fixed relative amplitude, i.e. ratio of predicted and observed amplitudes of peaks. **D:** Histogram of relative amplitudes.

For each unit, we convolved its peri-stimulus time histogram (PSTH) with its spontaneous STA EEG (Fig. 5A). This procedure results in an average (across click stimuli) contribution of this individual cell to the EEG; in response to a single click an NM unit typically produces only 1 − 4 spikes. Furthermore, because waveforms did not exhibit adaptation (Fig. 2), we assumed that the STA-EEG contributions add up linearly. In summary, averaging single-spike EEG contributions across clicks is equivalent to averaging the spiking responses of an NM unit, resulting in the PSTH, and then convolving the PSTH with the STA EEG.

The predicted contribution of the NM exemplary unit (Fig. 5A, green) had a 162 nV peak amplitude. The contribution peak was aligned in time with the peak of wave II (*P*_2_) of the click-driven ABR response with a difference of 240 *µ*s. The click-driven ABR response had an amplitude of 47 *µ*V (Fig. 5A, yellow), and thus this NM unit contributed about 0.28 ± 0.02% to the ABR wave II amplitude.

Also for the population, we tested which of the ABR peaks would show the largest contribution from the NM neurons. The average peak of ABR wave II (P2) was closest to the predicted peak (P) of NM contributions to the ABR (“P2-P” in Fig. 5B), with a median (± SE) relative delay of only −300 ± 20 *µ*s (*N* = 38 considering all stimulus levels, see also Table 3). However, the distribution of these relative P2-P delays was significantly different from the distribution of closest possible relative delays (“Pc-P” in Fig. 5B, mean: −250 ± 11 *µ*s, median = −160 *µ*s, 2-population t-test, Šidák-corrected for multiple comparisons: *p* = 0.011). Still, most of the closest delays stemmed from the wave II peak (23 out of 38), and a minority from the wave III peak (15 out of 38), with no significant difference in the stimulus levels between these groups (2-population t-test, *p* = 0.12). In contrast, the distributions of the relative delays P1-P and P3-P (medians: −950 ± 30 *µ*s and 710 ± 20 *µ*s, respectively) were both highly significantly different from the distribution of closest possible relative delays (*p <* 0.0001, 2-population t-tests).

**Table 3:**
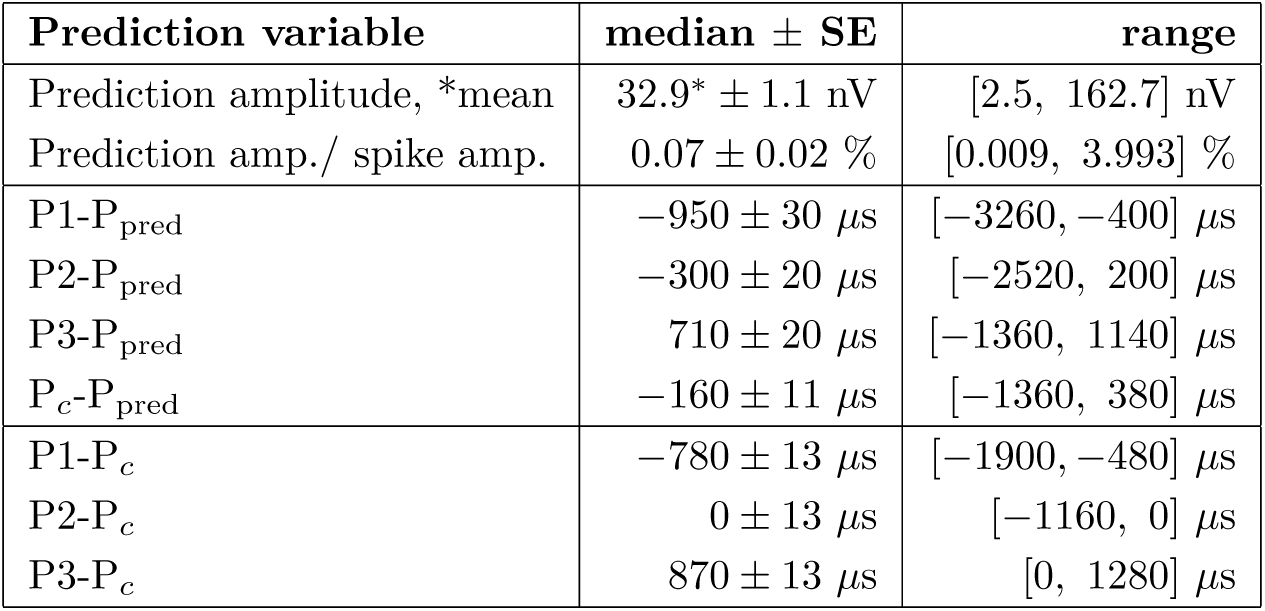
Prediction amplitudes and relative prediction delays of NM units. *N* = 38 predictions.

| Prediction variable | median ± SE | range |
| --- | --- | --- |
| Prediction amplitude, *mean | $32.9^* \pm 1.1$ nV | [2.5, 162.7] nV |
| Prediction amp./ spike amp. | $0.07 \pm 0.02$ % | [0.009, 3.993] % |
| P1-P<sub>pred</sub> | $-950 \pm 30$ μs | [-3260, -400] μs |
| P2-P<sub>pred</sub> | $-300 \pm 20$ μs | [-2520, 200] μs |
| P3-P<sub>pred</sub> | $710 \pm 20$ μs | [-1360, 1140] μs |
| P<sub>c</sub>-P<sub>pred</sub> | $-160 \pm 11$ μs | [-1360, 380] μs |
| P1-P<sub>c</sub> | $-780 \pm 13$ μs | [-1900, -480] μs |
| P2-P<sub>c</sub> | $0 \pm 13$ μs | [-1160, 0] μs |
| P3-P<sub>c</sub> | $870 \pm 13$ μs | [0, 1280] μs |

We previously showed that the level dependence was strong both for the ABR peak delays (Fig. 3B) and for the single-cell click-response delays (Fig. 2B), and that at the population level the slopes were indistinguishable. However, these slopes are insufficient to establish that at the single-cell level the relative timing between the prediction and the ABR peak(s) is level-independent. For example, the peak (but not the onset latency) of the PSTH will dominate the timing of the contribution peak. We therefore performed an N-way ANOVA based on the hypothesis that the delay of the ABR wave II peak and the delay of the contribution peak (both: *N* = 38) originated from the same level-dependent regression model. The group identity had no significant effect on the fit (*F*_1,69_ = 1.14, *p* = 0.35), whereas the click level did (*F*_5,69_ = 6.92, *p* = 0.011), indicating that there was no significant difference between the delay of the ABR wave II peak and the delay of the contribution peak. Furthermore there was no significant correlation between the level and the relative prediction delay with respect to the wave II peak delay (*p* = 0.35, *N* = 38; see Fig. 5B group P2-P). Such a level independence, in addition to the large spread of the relative delays in population, thus means that the ABR wave II peak delay cannot be predicted reliably by a single NM unit. Instead, wave II is expected to arise only when averaging over a large population of such predictions, as the units are all synchronized with the stimulus onset.

### Amplitudes of the predicted NM contributions were unexpectedly large

We predicted the average contribution of a putative single NM cell to the ABR by convolving the STA EEG with the cell’s click-evoked PSTH. Our hypothesis was that the contribution of a single cell to the ABR should be small because the ABR is a summation of most cells’ activities in the auditory brainstem. The amplitudes of the predicted NM single-cell contributions were broadly distributed (‘contribution peak’ in Fig. 5C). The mean predicted amplitude was 32.9 ± 1.1 nV (range 2.5 − 162.7 nV; Table 3), which is about half the average amplitude of STA EEGs (Table 2); this reduction of the predicted amplitude is the expected result of the convolution of the STA EEG with the PSTH. The relative amplitudes of the predictions ranged from 0.01% to 1% of the ABR wave II peak amplitudes, with a median (± SE) of 0.07 ± 0.02 % (Fig. 5C, D). One outlier (about 4%) was attributed to an unusually small ABR wave II peak amplitude. Neither the absolute amplitudes of the predictions (in nV), nor their relative amplitudes (in %) were significantly dependent on the stimulus level as a population (Pearson correlation coefficients: *CC* = 0.14, *p* = 0.40 and *CC* = −0.07, *p* = 0.64, respectively). However, the predicted amplitudes of individual cells were significantly dependent on stimulus level for 9 out of 16 NM units, when considering the logarithms of both ‘contribution peaks’ and the ‘ABR wave II peaks’, and using NM units’ identity as a random effect (GLM: *F*_17,21_ = 43, *p* = 2 · 10^−12^). All in all, stimulus level was not a good predictor of a unit’s relative contribution to the ABR wave II amplitude (Pearson correlation coefficient: 0.25, *p* = 0.13), even when using individual owls as a random effect (GLM: *F*_17,21_ = 1.69, *p* = 0.13). Thus, the relative peak amplitude of a given single unit to the ABR stayed approximately constant across stimulus levels.

As explained above, the amplitudes of contributions of single NM neurons to the amplitude of wave II of the ABR were large (about 0.1% i.e. 1*/*1, 000), which is unexpected when compared to the total number of neurons in NM of about 26, 000 (Han et al., 2024). One possibility is that NM STA EEG waveforms are generated not by a single NM neuron but by many structurally connected neurons in the auditory brain stem. In what follows we provide evidence against this idea.

To estimate how many units could contribute to the measured STA EEG of an NM unit, let us consider the NM circuit. An NM neuron is driven by AN endbulb synapses onto its cell body (for example, like the calyx of Held); these AN endbulb synapses are its only excitatory inputs. A single AN fiber is connected to 3 − 6 NM cells (Carr and Boudreau, 1991), and a spike in an AN fiber generates reliable response spikes in all connected NM cells (Brenowitz and Trussell, 2001; Wang et al., 2010). On the other hand, each NM cell receives inputs from 1 − 4 different AN fibers (Carr and Boudreau, 1991) and the spontaneous STA EEG of an NM cell could include contributions from them. Furthermore, there are contributions of 2 − 20 structurally connected NM cells in total: each of the 1 − 4 AN fibers project also to 2 − 5 other NM cells, which could all potentially contribute to the STA EEG of a single NM unit. In what follows, we neglect possible contributions from nucleus laminaris (NL) neurons that NM cells project to because these contributions appear with a delay of at least 1.3 ms after the NM activity (Wagner et al., 2005; McColgan et al., 2014).

We first focus on the contribution of AN fibers to the STA EEG of an NM unit. ABR wave I is assumed to be generated by AN activity (Melcher and Kiang, 1996), and we have presented evidence that wave II is generated by NM. Because the AN projects to NM and because ABR waves I and II have similarly large amplitudes (Fig. 3A), a contribution of connected AN units to the STA EEG of an NM unit seemed possible. This potential peak due to AN would be expected in advance of the STA EEG peak by more than the average synaptic delay of 0.51 ms (Table 1), but not more than 1.5 ms, considering the typical peak I-to-peak II inter-peak-intervals in the ABR of 0.5 − 1 ms. This range of delays is consistent with the (best-frequency dependent) length of the AN fibers from the basilar papilla to NM of ≈ 6 − 16 mm (Carr and Boudreau, 1991; Köppl, 1997a) and their predicted conduction velocities of 10 − 30 m/s (Carr and Boudreau, 1991; Köppl, 1997b; Seidl et al., 2010; McColgan et al., 2014). However, such an AN-related peak was not present in the grand average STA EEG of the NM units (Fig. 4C, bottom), and “peaks” were only occasionally observed in the STA EEG waveforms of individual NM units (colored parts of waveforms in Fig. 4C indicate significance). Somewhat surprisingly, but consistent with what we observed in the data, the AN contributions should typically be insignificant in the STA EEG of an NM unit because of two factors: Firstly, the activation of a single AN unit is sufficient to elicit a spike in an NM unit. Thus, an AN fiber participates only in a fraction (proportional to its spontaneous rate with respect to the typically higher spontaneous rate of the NM cell) of spikes of each NM unit. This leads to a reduction of each of the STA EEG waveforms of the participating AN units. Secondly, the variability in the structural conduction delays and synaptic delays between participating AN units and the NM unit reduces the summed contribution of the AN units. The variability of these conduction delays (jitter) is expected to be in the range of hundreds of microseconds because the axonal lengths can vary by as much as one millimeter between the branches of a single AN unit (Carr and Boudreau, 1991); furthermore, the delays that we measured between the prepotential and the NM spike ranged from 340 to 820 *µ*s (Table 1). Because this jitter is similar to or larger than the expected widths of the STA EEG waveforms of the 1 − 4 participating AN units (with widths presumably smaller than the width of ABR wave I), the amplitude of their average is much smaller than the sum of their amplitudes (Teleńczuk et al., 2015, their Fig. 4). Together, these two factors suffice to make the AN contribution insignificant in the grand average STA EEG of the NM units and in most of the single-unit STA EEG waveforms. The few cases in Fig. 4C where significant parts of waveforms could be observed at time lags *<* −0.5 ms might correspond to cases in which the NM unit gets input from a lower-than-average number of AN fibers (note that presumed NM units with spontaneous rates similarly low as AN units were excluded, see Fig. 1D).

We now turn to the contributions of structurally connected NM neurons (called “other units”) to the STA EEG of the recorded NM neuron (“unit of interest”). It is not possible to temporally separate the contributions of the other units from the contribution of the unit of interest because the expected average delays are identical. However, the same two effects for averaging across the other NM units (as argued above for AN units) apply here too: Firstly, the other NM cells participate only in a fraction of the spikes of our NM unit-of-interest, because the simultaneous spikes between the NM units need to be elicited by a mutually connected AN unit. This reduces the contribution of other NM units to the STA EEG; other, non-simultaneous spikes of other units are averaged out. Secondly, the variability in the structural conduction delays to the other NM units reduces their summed contribution to the STA EEG of the unit of interest. We expect that this variability is at least as large as the variability of AN-to-NM delays, which were already estimated to be in the range of several hundreds of microseconds. The variability of NM-to-NM delays may be even larger than the variability of the AN-to-NM connections because an NM-to-NM delay is the difference between two AN-to-NM delays (twofold increase in variance for independent AN-to-NM delays). In comparison, the widths of the STA EEG waveforms of the NM units are in the same range or even narrower (410 ± 9 *µ*s min-to-min around the largest significant peak, Table 2). Therefore, the summed contributions of the other NM cells should be markedly reduced compared to the STA EEG of the NM cell of interest (Teleńczuk et al., 2015, their Fig. 4). Together, these two effects are expected to greatly reduce the contributions of other NM cells to the STA EEG of the NM cell of interest. We therefore expect that the amplitudes of the STA EEGs of NM units in our data are similar or at least in the same order of magnitude as the amplitudes of “true” single STA EEGs of isolated NM cells (i.e., without other structurally connected NM cells).

## Discussion

Simultaneous recordings of ABRs and single units in barn owl NM demonstrated that individual spikes can make detectable contributions to the EEG, with a mean amplitude of 76 ± 4 nV (range 25 − 267 nV). The median single-unit contribution to the click-driven ABR was ≈ 0.1% of the elicited ABR wave II peak.

The time lag of the peak of the single-cell spike-triggered average (STA) EEG typically coincided with the rising phase of the extracellular NM spike waveform (−95 ± 12 *µ*s). However, the range of time lags was large (from −300 to +110 *µ*s excluding one outlier, Fig. 4D). This variability of time lags could be due to the variable position of the intracranial electrode: the time of the peak of the STA EEG is locked to the time of the spike initiation, but the propagation of the spike along the axon from the initiation site to the location of the intracranial recording electrode adds a variable delay. The longer this delay, the more negative the ‘time lag’. Furthermore, NM neurons have a variable spatial orientation, and this variable dipole axis can add variability to the time lag of the peak of the STA EEG. In contrast to the often negative time lag and the large variability we found, Teleńczuk et al. (2010) reported cortical STA EEGs for which the peak either coincided with the spike peak time, or for which the STA EEG had a rising phase at the spike time; and there was only a 100 *µ*s-range delay between the peaks. The grand average peak had some 50 − 100 *µ*s positive delay with respect to the spike peak. This may be explained by intracranial electrodes always being close to the soma and a preferred orientation of the dipole of pyramidal cells.

Let us compare the magnitude of obtained STA EEGs with those in other systems. We estimated the dipole moment *Q* of a spike generated by an NM cell based on the STA EEG peak potential *V*_STA_ with the dipole approximation (Malmivuo and Plonsey, 1995)

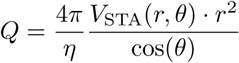

with constant tissue resistivity *η* = 2.47 Ωm (Logothetis et al., 2007), angle *θ* with respect to the dipole axis, and distance *r* of the EEG electrode from the source. The average intracranial recording depth below the dura was 10.2 ± 0.7 mm (mean ± SD). The active EEG electrode was positioned in the bone at ≈ 1 − 2 mm above the dura and ≈ 5 mm away from the intracranial electrode, which leads to *r* ≈ 12 mm. Furthermore, we assumed *θ* = 0 for the active EEG electrode. Thus, for the range of our STA EEG peak amplitudes (25 − 267 nV), the dipole moments range ≈ 20 − 200 nA mm. These dipole moments are larger than the dipole moments reported for cortical pyramidal neurons: Murakami and Okada (2006) found *Q* = 0.78 − 2.97 nA mm, which matches to data from pyramidal neurons of macaque monkeys (Teleńczuk et al., 2010) as well as to modeling results for rat and human cortical neurons (Næss et al., 2021). One reason for the inferred large dipole moments of NM neurons might be that the dipole approximation is not sufficient, i.e., our EEG electrodes were too close to the NM neuron and multipole moments of the electric field contributed to the measured potential.

The estimated dipole moment for NM spikes depends on the (unknown) spatial orientation of the dipole. Furthermore, the dipole moment depends on cell morphology (e.g. Næss et al., 2021), including the turns of the axon (Stegeman et al., 1987; Jewett et al., 1990), distribution of synaptic inputs (Gold et al., 2006; Lindén et al., 2010), spike generation site (Telenczuk et al., 2017), after-hyperpolarizing currents (Storm, 1987), and back-propagation of the spike (Gold et al., 2006; Telenczuk et al., 2017). NM neurons provide a useful contribution to the analysis of dipole moments because they typically have few or no dendrites (Carr and Boudreau, 1991), long (≈ 2 mm) and directed axons toward NL (McColgan et al., 2014), and putatively low input resistance of about 10 MΩ, similar to chicken NM (Kuba et al., 2015) and owl NL (Funabiki et al., 2011) neurons, all of which differentiates them, for example, from cortical neurons with typically higher input resistance and shorter and more dispersed axons. NM morphology led us to conclude that the STA EEG in the NM cells should originate from spiking in the cell body and the axon rather than from the synaptic and dendritic currents assumed for most cortical neurons (Mazzoni et al., 2010; Næss et al., 2021). The need for better understanding of the differences in the contributions of different types of neuronal morphologies to EEG calls for further modeling studies.

We predicted the average contribution of a single NM cell to the ABR by convolving the STA EEG with the cell’s click-evoked PSTH. The largest predicted single-cell contributions were ≈ 1% of the min-max amplitude of ABR wave II, and the median was ≈ 0.1% (Fig. 5C). Such large contributions were unexpected because NM has around 26, 000 neurons (Han et al., 2024) and most are activated by a click stimulus. There are several potential causes for this difference: even though the peaks of the predicted contributions of individual NM neurons aligned best with wave II of the ABR, those peaks showed temporal jitter (from −2.5 to +0.2 ms, Table 3, “P2-P”), which reduces the amplitude of the peak of the summed (across many NM neurons) ABR. Some units even made a negative contribution to the peak II. We also selected statistically significant STA EEGs, which likely biased measured amplitudes to large values. Finally, we might slightly overestimate the STA EEG of a single neuron due to correlated firing because of sparse structural correlations (see Results). Furthermore, temporal correlations in the spontaneous AN population activity, driven, for example, by body noises, are difficult to exclude.

To estimate the (click evoked) contribution of an NM cell to the ABR, we assumed that contributions of (possibly several) spikes of the NM cell add up, and we therefore convolved the STA EEG with the PSTH (Fig. 5A). The underlying linear-summation assumption is reasonable because there is little adaptation of spike waveforms (e.g. Figs. 1B and 2A), and AN-to-NM synapses in chicken show little or no adaptation within the first few driven spikes (Avissar et al., 2007; Ahn and MacLeod, 2016). Furthermore, possibly non-linear inhibitory feedback from the superior olivary nucleus (SON) should not contribute during click onset (Lachica et al., 1994; Monsivais et al., 2000; Coleman et al., 2011).

The compound effect of a neuronal population to the ABR depends on the synchronization of the cells within the population (Kuokkanen et al., 2010; Ahlfors et al., 2010a,b; Lindén et al., 2011). Temporal synchrony is famously precise in the auditory brainstem (Kiang, 1965) leading even to signals that can be recorded at the scalp more than a centimeter from their source (McColgan et al., 2017). Note that the ABR, exhibiting several waves, is a sum of several subsequently activated neural populations. Thus, assumptions of the populations’ spatial alignment and temporal synchronization underlie, at least implicitly, all ABR models (Melcher and Kiang, 1996; Ungan et al., 1997; Dau, 2003; Goksoy et al., 2005; Riedel and Kollmeier, 2006; Colburn et al., 2008; Schaette and McAlpine, 2011; Rønne et al., 2012; Verhulst et al., 2015, 2018). Our results suggest that NM responses alone are sufficient to produce wave II, but a thorough quantification would require additional modeling to consider the variable geometry of NM cells. Furthermore, other sources, such as nucleus angularis (NA) (Takahashi and Konishi, 1988; Köppl and Carr, 2003) likely contribute to wave II. NA, like NM, is a first-order auditory nucleus with similar average onset latencies as NM (Köppl and Carr, 2003), and its contributions are expected to be temporally aligned with the ABR wave II. However, the observed variation in onset latencies (≈ 1.5 − 4.5 ms for 20 − 35 dB tones, Köppl and Carr, 2003) and between response types in NA raises questions about their coherence in generating a collective ABR peak (Sachs and Sinnott, 1978; Soares et al., 2002; Köppl and Carr, 2003).

Other brainstem structures, such as NL and SON, can be excluded as wave II sources because they have longer response latencies than NM (Lachica et al., 1994; Yang et al., 1999; Monsivais et al., 2000; Burger et al., 2005). McColgan et al. (2017) estimated that the low-frequency field (*<* 1 kHz) created by branching patterns of the NM axons in NL could collectively contribute microvolt excursions in the scalp EEG recordings. This contribution is expected to be more aligned with ABR peak III than peak II, considering a conduction delay of at least 1.3 ms between the NM cell body response and the responses from their axonal arbors in the NL (Carr and Konishi, 1990; Köppl, 1997c; Wagner et al., 2005; McColgan et al., 2014). The amplitude of high-frequency (*>* 1 kHz) oscillations generated in NL (‘neurophonic’) decays rapidly outside NL (Carr et al., 2015; McColgan et al., 2017) and is not expected to contribute to the EEG. Finally, understanding potential contributions of the AN fiber branching pattern onto NM neurons to ABR wave II should require further modeling and experimentation.

There are clear differences between the *unitary response* (UR), used in ABR modeling, and the STA EEG (and its convolution with the PSTH) that we have measured, despite the UR being defined as the expected average spike-triggered response of a single neuronal source at the EEG electrode. For one, the UR, as often used in ABR models, is typically derived from the temporally correlated driven responses (ABRs) by deconvolution, and includes also the structurally correlated cascade of activation of any neuronal sources associated with the spike in a single auditory nerve fiber (Dau, 2003; Rønne et al., 2012; Verhulst et al., 2015, 2018). By contrast, we tried to minimize such correlations in our STA EEG by using spontaneous spikes, and show only the scalp contribution of single NM cells. Secondly, the UR has the same average waveform for all sources, disregarding any variation in the neuron population or even between neuron types. By contrast, our STA EEGs include the large variability present in the NM cell population. Defining the STA EEG for a group of single neurons in a single nucleus should help limit the number of possible realistic URs. Given the wide range of the STA EEG responses, our data suggest that it is unlikely that a single NM spike-triggered average EEG waveform represents the UR. Instead, an NM UR can be derived from the sum of the STA EEG responses.

## Acknowledgments

We thank Go Ashida, Ghadi El Hasbani, Tizia Kaplan, Lutz Kettler and Nadine Thiele for helpful discussions, and thank Waisudin Kamal for his assistance with cell counts.

Supported by NSF CRCNS IOS1516357, by the National Institute on Deafness and Other Communications Disorders (NIDCD DC00436 and DC019341), and the Bundesministerium für Bildung und Forschung (BMBF): German – US-American collaboration “Field Potentials in the Auditory System” as part of the NSF/NIH/ANR/BMBF/BSF CRCNS program, 01GQ1505A and 01GQ1505B. The research was further funded by the Deutsche Forschungsgemeinschaft (DFG, German Research Foundation) grant nr. 502188599.

